# Rate Dependent Cochlear Outer Hair Cell Force Generation: Models and Parameter Estimation

**DOI:** 10.1101/2023.12.13.571371

**Authors:** Wen Cai, Karl Grosh

**Author notes:** Contributing authors. These authors contributed equally to this work.

## Abstract

The outer hair cells (OHCs) of the mammalian cochlea are the mediators of an active, nonlinear electromechanical process necessary for sensitive, frequency specific hearing. The membrane protein prestin conveys to the OHC a piezoelectric-like behavior hypothesized to actuate a high frequency, cycle-by-cycle conversion of electrical to mechanical energy to boost cochlear responses to low-level sound. This hypothesis has been debated for decades, and we address two key remaining issues: the influence of the rate dependence of conformal changes in prestin and the OHC transmembrane impedance. We develop a theoretical electromechanical model of the OHC that explicitly includes rate dependence of conformal transitions, viscoelasticity, and piezoelectricity. Using this theory, we show the influence of rate dependence and viscoelasticity on electromechanical force generation. Further, we stress the importance of using the correct mechanical boundary conditions when estimating the transmembrane capacitance. Finally, a set of experiments is described to uniquely estimate the constitutive properties of the OHC from whole-cell measurements.

## 1 Introduction

Mammalian hearing relies on a biologically vulnerable, active process in the cochlea that is responsible for spatially-dependent tuning and subnanometer threshold responses to sound. The active process produces sharp tuning and high gain at low sound levels, features that are dramatically reduced when sound pressure levels are increased. Two key properties of cochlear outer hair cells (OHCs) are central to this process. First, the apical pole of each OHC has ∼ 100 stereocilia arranged in three rows, with two rows equipped with mechanoelectrical-transducer (MET) channels whose conductance depends on acoustically-induced stereocilia motion [1]. This variable conductance modulates the resting potential of the cell generating a time-varying MET current (*I*_*met*_) (e.g., 2). Second, the lateral membrane of the OHC is densely packed with the transmembrane protein prestin [3]. Because of the unique constitutive properties of prestin, the OHC is capable of converting mechanical stimulation into an electrical response (forward transduction) and electrical stimulation into a mechanical response (reverse transduction), a feature known as somatic motility [4]. The electromechanics of the stereocilia and soma have been implicated as the mediator of the active process necessary for sensitive hearing [1, 4]. Experimentally, any genetic, pharmacological, or mechanical intervention that degrades either OHC MET channel or prestin functionality also disrupts and often eliminates this enhancement, rendering hearing insensitive, broadly tuned, and largely linear in its response (e.g., 1, 5–7).

Functionally, the somatic-based cochlear amplifier (CA) theory couples MET currents and somatic motility in a feedback mechanism where the resting endocochlear potential powers an electrically generated, amplifying somatic force [4]. This theory, which has been proposed ever since OHC electromotility was first identified [8], can be used to explain key aspects of the cochlear function [9, 10], and has generated the most support in the field [1]. However, for the somatic-based CA hypothesis to be valid, prestin-mediated forcing must be viable at acoustic frequencies, up to 100 kHz, and high frequency forcing is challenged mainly on two grounds. First, the speed of conformal changes of prestin may limit the upper frequency of functionality for somatic motility [11]. Second, if basolateral membrane capacitance is too large, then the MET current-induced transmembrane voltage may be too small to generate sufficient somatic force to drive the active feedback process [12]. This concern arises because the OHC operates at frequencies where capacitance, not resistance, dominates the basolateral membrane impedance [13–15], even under the estimates of conductance and capacitance most favorable to high-frequency amplification [2]. Due to these over 30-year-old challenges, other theories of cochlear enhancement continue to be actively pursued [16–18], and it has been speculated the somatic-based CA may work cooperatively with other modes of enhancement, or operate in different regions along the cochlear spiral (e.g., 4).

The purpose of this paper is to present a set of theoretical results to facilitate a critical investigation of somatic-motility based CA theory. Recently, OHC constitutive theory has been shown to match experimental results nicely [19]. However, the model is overfit since insufficient experimental data exist to uniquely identify all parameters. Because of the overfitting, assumptions regarding parameters are required to finalize a model, leading to open questions about the effectiveness of somatic electromotile forces in amplifying sounds (e.g., 20, 21), even though there is significant *in vivo* and *in vitro* evidence supporting high frequency somatic motility (e.g., 9, 22–25). We seek to demonstrate precisely which parameters are needed for models, how estimates of these parameters might be obtained through experiments (with a special emphasis on mechanical boundary conditions), and the significance of this theory in testing hypotheses of cochlear enhancement. Through this analysis, we hope to move further along the path toward clarifying this longstanding debate.

## 2 Results

### OHC Constitutive Model

Somatic-based cochlear amplification is a multi-step process where sound-evoked vibrations of the stereocilia cause fluctuating MET currents at the stereocilia which in turn give rise to transmembrane potential fluctuations. These voltage fluctuations generate time-varying forces through reverse transduction in the OHC that, if timed correctly, enhance the mechanical response in a cycle-by-cycle fashion drawing power from the resting endocochlear electrical DC potential. This interaction creates an electromechanical feedback loop as the stereocilia motion initiates a chain of events that ultimately leads to voltage generated forces that affect stereocilia motion [4]. Although many factors influence the strength of this interaction, including global wave propagation [26], current spread [27], and the dynamics of the organ of Corti [14, 28], the axial force generated by the OHC is the key link in the theory. In the cochlea, a controlled voltage or force is not directly applied to the cells themselves. Rather, the response stems from a system-level coupling of the fluid, mechanical, and electrical domains stimulated by external acoustic stimulation [1, 29]. However, in order to determine the constitutive behavior of the cells, controlled *in vitro* experimental results provide the parameters for an isolated model that can then be inserted into a global cochlear model (e.g., 10, 30). Therefore, in this section, we develop a model for axial forces suitable for modeling *in vitro* experiments [31] and for inclusion into a global model of the cochlea [10, 30].

The OHC is a fluid-filled shell of nearly cylindrical shape *in situ* and when extracted from the organ of Corti [3, 31]. Under quiescent electrical and mechanical conditions, the cell is in equilibrium under a turgor pressure (*P*_0_), axial force (*T*_0_), and resting transmembrane voltage (*V*_0_) as pictured in Fig. 1(a). The cell wall is comprised of a densely packed mosaic of the transmembrane motor protein prestin sandwiched in the lipid bilayer [32, 33]. The plasma membrane is connected in parallel to the cortical lattice [3, 34], but the contribution of the cortical lattice to the OHC axial stiffness has been estimated to be much smaller than that of the plasma membrane [34, 35]; hence, we omit it here.

**Fig. 1.**
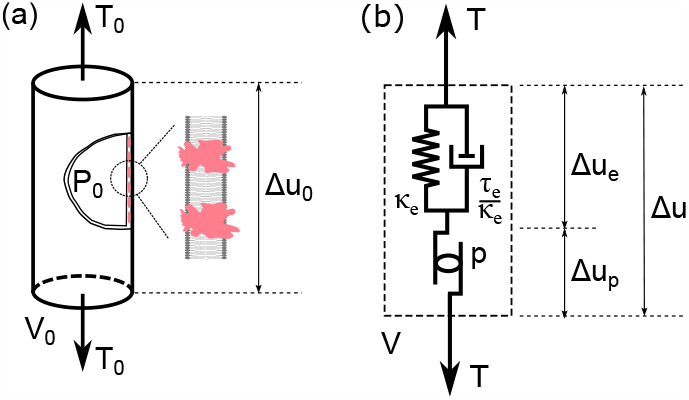
(a) Axially loaded OHC at equilibrium with the external force *T*_0_, resting potential *V*_0_, turgor pressure *P*_0_, and static deformation ∆*u*_0_. The inset shows a cartoon of the plasma membrane cross-section showing the lipid bilayer (in grey) with the prestin (in red) spanning the membrane [33]. (b) OHC model. Based on the structure of the plasma membrane, the OHC can be modeled as an electromotile element *p* representing the overall prestin contributions in series with a viscoelastic element characterized by compliance *κ*_*e*_ and a rate constant *τ*_*e*_ (or damping constant of *τ*_*e*_*/κ*_*e*_) [19]. The electromotile element *p* represents the nonlinear, and rate dependent mechanical and electrical function of the transmembrane proteins (see Methods section). The notation *δ* represents perturbations of the response quantities away from equilibrium so that the total deformation is given by ∆*u* = ∆*u*_0_ + *δu*, the prestin deformation by ∆*u*_*p*_ = ∆*u*_*p*0_ + *δu*_*p*_, viscoelastic deformation by ∆*u*_*e*_ = ∆*u*_*e*0_ + *δu*_*e*_, force by *T* = *T*_0_ + *δT*, voltage by *V* = *V*_0_ + *δV*.

Using the equations of equilibrium in the circumferential (or hoop) direction and the incompressibility constraint of the intracellular OHC fluid, the intra-OHC pressure changes and radial deformation can be eliminated from the axial load-deformation relations [31, 36], resulting in a model whose mechanical variables are represented by the axial deformation, ∆*u*, which can be seen as the combination of the net deformation of all motor elements ∆*u*_*p*_ and the lipid bilayer ∆*u*_*e*_ in Fig. 1(b) [19, 37, 38]. Similarly, the transmembrane charge (*Q*) is additively decomposed into the nonlinear charge associated with the motor process (*Q*_*nl*_ = *Q*_*nl*0_ + *δQ*_*nl*_), and the linear charge component (*Q*_*lin*_) as these components are electrically in parallel and experience the transmembrane voltage *V*. The linear contribution *Q*_*lin*_ is often carefully identified and subtracted from experimentally presented results (e.g., 39).

Since the perturbation of the displacement and voltage away from equilibrium *in vivo* are typically small (e.g., the receptor potential changes are at most 10 *mV* for most mammalian auditory systems [40]), we develop the linearized relations about equilibrium to investigate the somatic mechanics of the OHC. In the Methods section, we derive the coupled, time-dependent differential equations relating the small perturbations of the total OHC displacement, *δu*, and nonlinear charge, *δQ*_*nl*_, caused by small perturbations of the force, *δT*, and transmembrane voltage, *δV*. Further, we transform these equations from the time domain into the frequency domain (using *e*^*jωt*^ time dependence, e.g., *δu = ũ e*^*jωt*^), where the diacritical tilde indicates the complex representation of the frequency dependence for each response variable (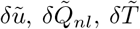, and 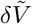). As derived in the Methods section, Eqs. 41-42, the frequency domain relations in displacement-charge form are given by

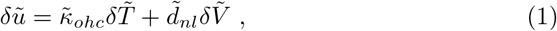

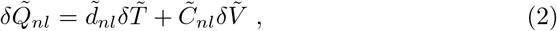

and rearranged in force-charge form as

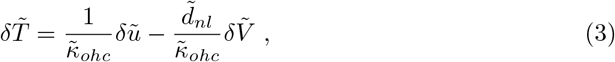

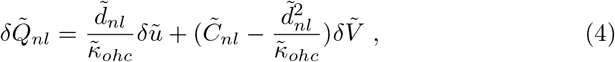

where

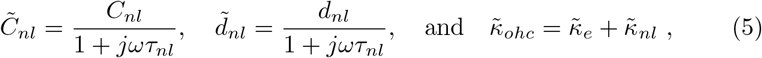

with

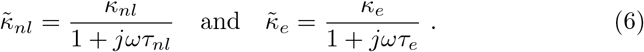

In the linearized relations, the material properties with the subscript *nl* are dependent upon the resting state, including the quasi-static compliance *κ*_*nl*_, piezoelectric strain coefficient *d*_*nl*_, and free nonlinear capacitance (NLC), *C*_*nl*_ [21, 31, 38, 41]. The time constant associated with conformal changes of prestin is *τ*_*nl*_. According to the absolute rate Eyring theory (see Methods section, Eq. 31 and 21), *τ*_*nl*_ attains its maximum value when the motor is at the half-probability state and decreases about that point. The viscous damping in the elastic domain is characterized by the compliance of the lipid bilayer *κ*_*e*_ and the rate constant *τ*_*e*_. The OHC material parameters adorned with a tilde are complex and frequency dependent (as defined in Eqs. 5-6) while the parameters without a tilde represent the values measured under quasi-static (low frequency) conditions. The total OHC compliance is given by 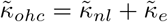.

Equations 1-4 represent the generalized constitutive behavior of the basolateral wall of the OHC. The boundary conditions (electrical or mechanical) must be prescribed in order to make predictions. Both the displacement-charge and the force-charge forms of the equations are useful for interpreting experiments. In modeling, the force-charge form is the most useful as it forms the basis for the inclusion of somatic OHC electromechanics to both the MET currents and to the dynamics of the organ of Corti [14, 30, 42]. We close this section with three remarks relating to the force-charge form of the equations.

#### Remark 1

The model predicts that electromechanical force generation by the transmembrane potential is not always low-pass filtered, and has a different frequency behavior than either the piezoelectric strain coefficient 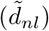 or load-free NLC 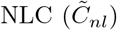. The electromechanical force 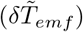 generated by the transmembrane potential in Eq. 3 is

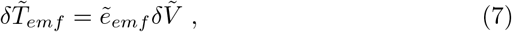

where the electromechanical force coefficient (sometimes called the isometric force factor [31]) is

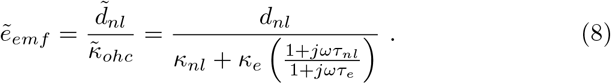

While both 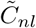 and 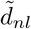 depend on a constant divided by (1 + *jωτ*_*nl*_) (see Eq. 5), the frequency dependence of 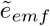 is slightly more complicated, and the low and high frequency asymptotes are given by

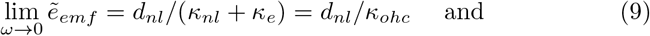

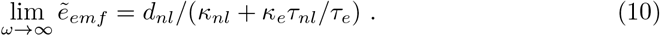

The frequency dependence of 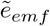 depends on the ratios of *κ*_*e*_*/κ*_*nl*_ and *τ*_*e*_*/τ*_*nl*_. Two special cases are notable. First, when *τ*_*e*_ = 0, the 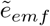 is low pass filtered, and second, when 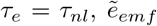 is real-valued and frequency independent. Finally, since both the compliance (*κ*_*ohc*_) and piezoelectric strain coefficient (*d*_*nl*_) scale with length [11, 22, 31], Eq. 8 shows that *e*_*emf*_ = *d*_*nl*_*/κ*_*ohc*_ is expected to be roughly constant with OHC length, as found by 31.

#### Remark 2

The total basolateral electrical impedance, 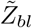, for use in a force-charge OHC circuit model [14] is given by

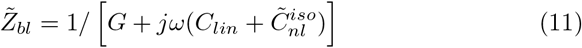

where *G* is the conductance, *C*_*lin*_ is the linear capacitance, and 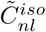is the complex, frequency-dependent isometric NLC, found from Eq. 4 when 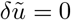 as:

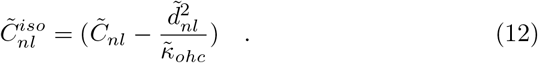

Both *G* and 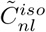 are functions of the resting potential [2], while *C*_*lin*_ is assumed to be constant [43]. As frequency increases, 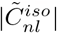 decreases as |1*/*(1 + *jωτ*_*nl*_) |, thereby increasing the overall transmembrane impedance. The increased lateral impedance would also increase the transmembrane potential due to MET current, a phenomenon that could impact electrical-to-mechanical force production. Finally, it has not been widely recognized that it is the isometric capacitance, 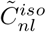, that should be used in the force-displacement version of the OHC constitutive relations, Eq. 4.

#### Remark 3

The so-called RC filter frequency is set by the combination of the isometric NLC, the basolateral conductance, and linear capacitance, as given Eq. 11. Although direct estimates of the 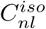 for a cylindrical, non-trypsinized cell do not appear to be available, using estimates of *d*_*nl*_ ≈ 20 nm/mV [44] and *κ*_*ohc*_ ≈ 100 m/N [31] for a 50-micron OHC yield 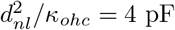 resulting a 25% reduction of the peak *C*_*nl*_ (∼ 16 pF [43]). If we restrict our constitutive theory such that nonlinear deformation *u*_*p*_ and nonlinear charge *Q*_*nl*_ are linearly related (see Methods section, Eqs. 22, 23, and 34 and 19, 36, 45), then 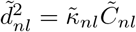 which further yields:

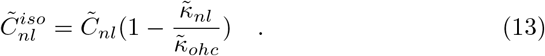

Using the above parameters, we would expect 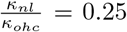, indicating that most of the OHC compliance stems from the cell membrane. We can also see that 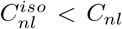. This places an imperative on experimentally determining the ratios (either 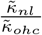 or 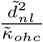), to estimate the frequency dependence of model parameters effecting 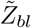.

### Mechanically Loaded OHC Model

Eqs. 1-4 embody the intrinsic dynamic constitutive model equations for the OHC. These relations not only provide a mechanistic description of OHC function from isolated measurements, but also can be incorporated into a model of the cochlea. Before doing so, experimental measurements of the underlying material properties are needed. Naturally, any experimental configuration used to estimate the OHC properties will influence the response. We consider a simple model where a location near the basal pole of the OHC is fixed and the apical pole is loaded by a spring-mass-damper system meant to approximately model the interaction with the surrounding media (e.g., loading by the exterior fluid and flexible fiber), as shown in Fig. 2. The external load can be represented by a dynamic stiffness term

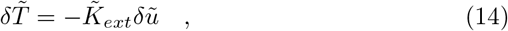

where

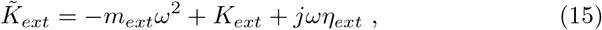

and *K*_*ext*_ is the external stiffness, *m*_*ext*_ is the external mass loading the OHC, *η*_*ext*_ is the external damping. Using Eq. 14, 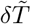 can be eliminated from Eq. 3. Solving for the resulting motion 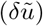 under known voltage sinusoidal voltage excitation 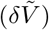, we find the frequency dependent transfer function between voltage and displacement, 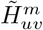 is given by

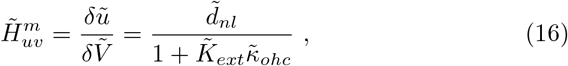

and this quantity is sometimes called electromotility (eM) in the literature. The measured NLC, 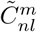, is given by

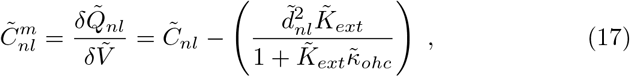

**Fig. 2.**
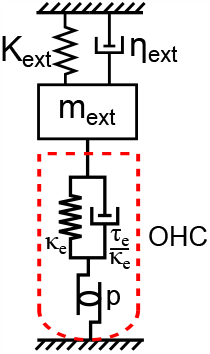
OHC connected to a load modeled as a spring-mass-damper system in order to approximately simulate the external environment. The stiffness *K*_*ext*_ is to simulate the effect of a flexible fiber [31] or atomic force microscope probe [22]. The mass *m*_*ext*_ and damping *η*_*ext*_ represent the influence of the entrained fluid mass [22] and viscous damping force from the external fluid medium [3, 19, 22, 46]. The basal pole is fixed.The external impedance can be written as 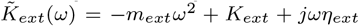. The basal pole (or some location nearby) is held fixed by the patch-clamp pipette attachment [44].

In both cases, the responses consist of not only the intrinsic OHC properties but also the extrinsic stiffness, damping, and mass effects. The extent of the influence of the external terms depends on their magnitude compared to the intrinsic properties. For instance, in the low frequency or quasi-static regime, the measured electrical capacitance, 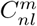, is an estimate of the free NLC, *C*_*nl*_ if *K*_*ext*_ → 0. If *K*_*ext*_ approaches infinity, the measured NLC estimates isometric NLC 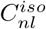.

For the loading condition shown in Fig. 2, under the voltage clamp condition, the power delivered to the external load by the OHC is the time average of the electromechanical force (Eq. 7) times the velocity over one cycle as given by:

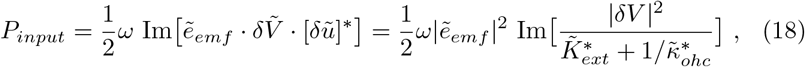

where []^∗^ indicates the complex conjugate, Im[] indicates the imaginary part of the complex number. The rate dependent electromechanical force coefficient 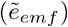 plays a central role in characterizing the power transfer as it did for forcing. Our expression for power differs from 19 because we consider the total work done by the OHC on the outside world rather than the available power from the motor. We reckon the power to the total displacement rather than just the prestin-related deformation term.

The measured eM and NLC under arbitrary mechanical loading do not directly yield the material properties as a function of frequency. Rather, experimental measurements must be processed to deliver the boundary– condition independent material properties used in Eqs. 1-6 because of the influence of external mechanical and electrical boundary conditions. One main finding is that even though the linearized value of the NLC, 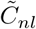, and piezoelectric strain factor, 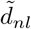, might demonstrate low-pass filtering, the electromechanical force coefficient, 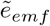, does not share that dependence. The key relationship between *τ*_*e*_ (the viscoelastic time constant) and *τ*_*nl*_ (the protein conformal time constant) defines if the force filtering falls off dramatically at higher frequencies, or maintains a nearly all-pass behavior (Eqs. 8-10), which has yet to be clarified.

### Model Predictions and Experimental Results

The six intrinsic OHC parameters: *κ*_*nl*_, *κ*_*e*_, *τ*_*nl*_, *τ*_*e*_, *C*_*nl*_, and *d*_*nl*_ along with the three external loading parameters define the response of the externally loaded the OHC under a small sinusoidal voltage fluctuation about equilibrium, as given in Eqs. 16 and 17. The model is fit to the experimental data from 22 and 47 with the parameters given in Table 1 as described in the Methods section.

**Table 1.** Parameter Values in the OHC Model.

| | $\kappa_{nl}$<br>[$\frac{m}{N}$] | $\kappa_e$<br>[$\frac{m}{N}$] | $\kappa_{ohc}$<br>[$\frac{m}{N}$] | $d_{nl}$<br>[$\frac{nm}{mV}$] | $C_{nl}$<br>[pF] | $\tau_{nl}$<br>[$\mu s$] | $\eta_{ext}$<br>[$\frac{N \cdot \mu s}{m}$] | $m_{ext}$<br>[pg] | length<br>[$\mu m$] |
| --- | --- | --- | --- | --- | --- | --- | --- | --- | --- |
| OHC 65 | 1.7 | 125.3 | 127 | -3.066 | — | 0.7 | 0.081 | 176.86 | 55.6 |
| OHC 84 | 0.32 | 74.08 | 74.4 | -0.575 | — | 0.1 | 0.055 | 76.34 | 24 |
| OHC 1 | 5.5 | 117.5 | 123 | -12.3 | 27.7 | 70 | 5 | 300 | 78 |

In Fig. 3, the simulated electromechanical responses are compared to measurements of 22 under two separate mechanical boundary conditions. The experiments were done with the isolated OHC partly inserted into a microchamber, and the simulated displacement is corrected to consider the whole cell configuration as in 22. First, the voltage-excited displacement responses 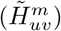 for two OHCs (OHC 65 and OHC 84) under fluid loading only (i.e., with no external stiffness applied) are considered. Both the magnitude (Fig. 3(a)) and phase (Fig. 3(b)) of the simulated OHC displacement responses match the experimental results well. The low pass nature of the response for this choice of model parameters is controlled by the OHC stiffness and the external fluid damping (*η*_*ext*_). The influence of *τ*_*e*_ is minor, only seen near 100 kHz in the phase response for OHC 65. In Figs. 3(c) and 3(d), the averaged experimental force output driven by sinusoidal voltage excitation is shown from three different OHCs (not OHC 65 and OHC 84) loaded by an atomic force microscope (AFM) probe with blue triangles [22]. The AFM probe is much stiffer than the OHC; hence, this provides an estimate of the electromechanical force coefficient 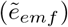. In Figs. 3(c) and 3(d), we compare these averaged experimental data to our model predictions using the parameters estimated for OHC 65 and OHC 84. The results of OHC 84 fall very close to the mean value of 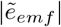 while the electromechanical force coefficient for OHC 64 is below this mean value. When the excitation frequency exceeds 50 kHz, variations in the experimental data were attributed to the resonance in the fixture [22].

**Fig. 3.**
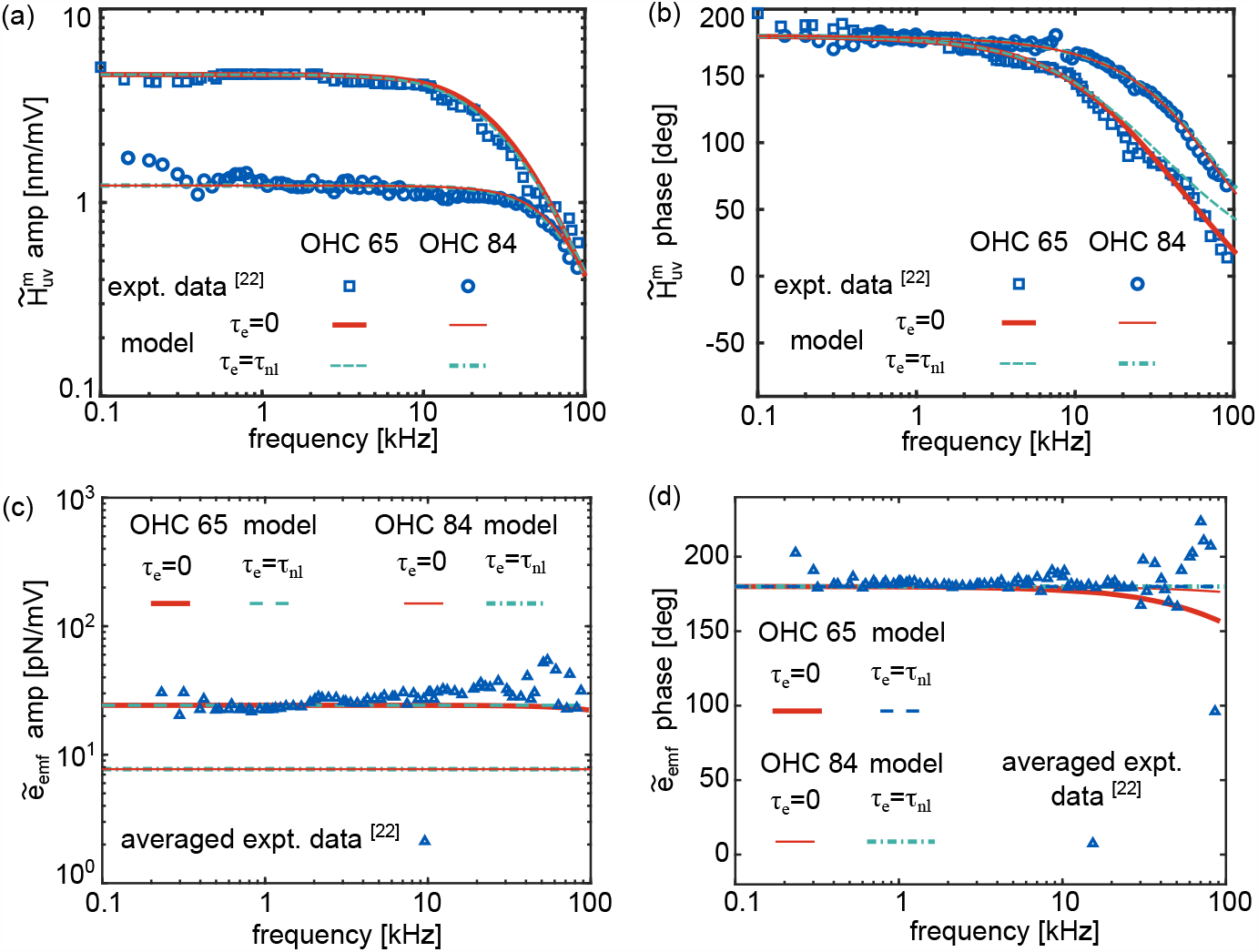
A comparison of theoretical predictions (solid lines) and experiments (symbols) of the (a) magnitude and (b) phase of OHC eM 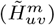 for OHC 65 and OHC 84 (the experimental data were taken from Figure 2 in 22). In (c) and (d) we present theoretical predictions of the magnitude (c) and phase (d) 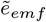 for parameters fit to OHC 65 and OHC 84 to averaged measured from three different cells (not OHC 65 and OHC 84) (experimental data are taken from Figure 4 in 22). The range of the magnitude of the measured electromechanical force coefficient for 15 cells reported in 22 is 3 to 53 pN/mV.

The experimental data in 22 are influential as they demonstrate that OHC high-frequency electromechanical force generation in an *in vitro* setting is viable up to and beyond acoustically relevant frequencies. This result is consistent with *in vivo* electrical stimulation results [24, 25]. However, both older (e.g., 11) and more recent experimental results (e.g., 48) based on NLC and eM (but not on force generation) indicate that the transition rate might be much slower than what is found in 22. One possible explanation for this difference is due to the resting potential dependence of *τ*_*nl*_, which is predicted to be slowest near *V*_*peak*_, the holding voltage of peak NLC or the 1/2 probability point [49]. It is therefore conjectured that since 22 likely used a holding voltage away from the 1/2 probability point, their results were non-physiologically fast. Theory (see Methods section Eq. 30) and experiment (see Fig. 6 of 21) support the state dependence of rate constants. However, even the fastest experimental rate estimated by 21 is a factor ∼ 5 slower than those seen in 22, indicating a gap in our understanding of the difference between these two measurements.

To explore if our model could also replicate results measured at *V*_*peak*_, we simulated data that were taken at a resting voltage of *V*_*peak*_. In Fig. 4(a), model predictions using parameters for OHC 1 from Table 1 while varying *τ*_*e*_ over three values are compared to the whole-cell patch clamped NLC and eM experimental data (data are taken from the middle panel of Fig. 4B in 47). The predicted NLC matches the data and is insensitive to the selection of *τ*_*e*_. The best fits of eM arise from choosing *τ*_*e*_ = *τ*_*nl*_*/*2, although all three choices fit well. For the parameters given in Table 1, the cut-off frequency of the NLC is governed by the rate *τ*_*nl*_, whereas the cut-off frequency of the eM is controlled by the external mass and viscous loading.

**Fig. 4.**
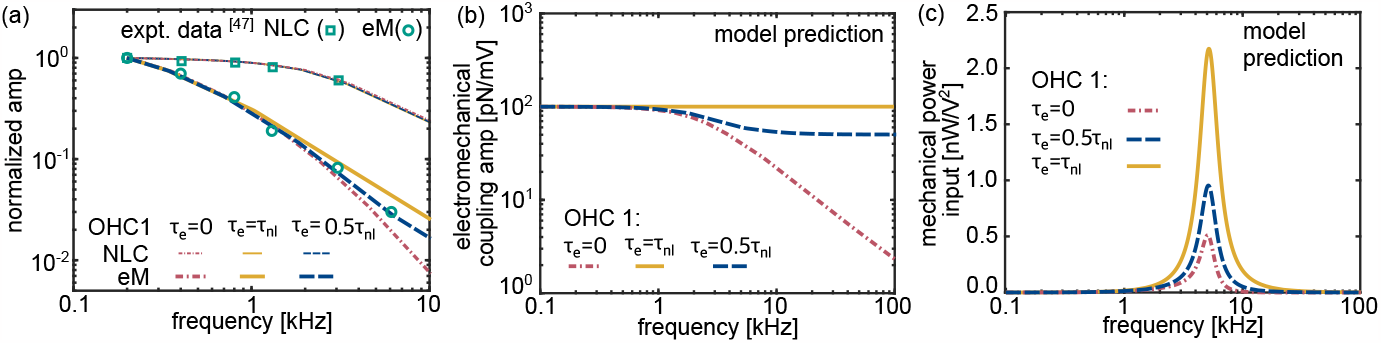
(a) Predicted non-dimensional amplitude of eM and NLC compared to the experimental data in figure 4B (middle panel) in 47 (b) Amplitude of the electromechanical force coefficient 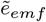from model prediction for OHC 1. (c) Electrical-to-mechanical power transfer to the impedance load in the frequency domain with different rate dependence, simulated with mass *m*_*ext*_ = 0.16*μg, K*_*ext*_ = 0.16*N/m*, and *η*_*ext*_ = 1.74*μN · s/m*. The OHC parameters are taken from OHC 1 in Table 1.

While the variation of *τ*_*e*_ does not alter the NLC responses and has a minor effect on eM, reducing *τ*_*e*_ has a dramatic effect on the electromechanical force coefficient 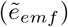, as shown in Fig. 4(b). When *τ*_*e*_ = 0, the force factor of OHC 1 acts as a low-pass filter, with a cut-off frequency of 2.44 kHz (red dot-dashed line); when *τ*_*e*_ = *τ*_*nl*_, the force factor generated from the OHC is constant through the whole frequency range (solid gold line), acting as an all-pass filter; and when *τ*_*e*_ is 0.5*τ*_*nl*_, a dual-plateau filtering is demonstrated for OHC 1 (blue dashed line). No experimental data for 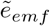 were available over the frequencies of interest.

In Fig. 4(c), the power delivered to an external load by OHC 1 is computed. In these simulations, the external load is selected to have a resonance of ≈5 kHz, more than twice the cut-off frequency of the cell predicted by 1*/*2*πτ*_*nl*_, by adding a mass *m*_*ext*_ = 0.16 *μg* and using a stiffness *K*_*ext*_ = 0.16 *N/m*. The external damping *η*_*ext*_ is also modified to be 1.74 *μN* · *s/m* to simulate an under-damped condition (*Q* ≈ 3) as seen in a healthy, active cochlea [9]. The power, *P*_*input*_, is plotted in Fig. 4(c) for three different values of *τ*_*e*_. The nano-Watt/*V* ^2^ input power in Fig. 4(c) for all cases is consistent with predictions in the literature [19, 50, 51]. As seen with 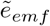, the value of the viscoelastic time constant *τ*_*e*_ influences *P*_*input*_. When *τ*_*e*_ = *τ*_*nl*_, the mechanical power input is enhanced by more than 4 times compared with the case that *τ*_*e*_ = 0, and twice that of *τ*_*e*_ = *τ*_*nl*_*/*2. We found that the *τ*_*nl*_ has a significant influence on the input power: keeping other factors the same and reducing *τ*_*nl*_ by half results in around a 3-fold enhancement in the power, while increasing *τ*_*nl*_ results in a decrease in power generation.

In summary, if 1*/*(2*πτ*_*nl*_) is commensurate with acoustic frequencies, then the values of *τ*_*e*_ and *τ*_*nl*_ play important roles in the electromechanical force coefficient of the OHC, as seen in Fig. 4(b) and (c). On the other hand, if both time constants are faster than relevant acoustic time scales, as in Fig. 3 (models of OHC 65 and OHC 84), then external loading dominates the rate-dependent phenomenon. While the models we present here can fit existing data, they are not uniquely constrained by experiment. The data from eM and NLC measurements alone are insufficient to determine all the parameters.

### Protocol for estimating OHC model parameters from experimental measurements

In order to characterize the ability of OHCs to actuate at high frequencies, the frequency-dependent magnitude and phase of 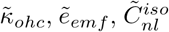 (see Eqs. 3-4) must be estimated from experiments. In this section, we put forth an experimental approach for obtaining the model parameters requiring only the quasi-static (low frequency) stiffness of any external probe to be quantified before the measurements.

Consider two separate experiments, performed under distinct mechanical boundary conditions. Ideally, in one experiment, the apical pole of the OHC is nearly free, loaded only by the external fluid, as in Fig. 5(a). In the other experiment, the OHC is constrained by a stiff fiber, as in Fig. 5(b). The rationale for using nearly load free and isometric conditions is to generate the largest difference between responses. In both experiments, the patch is preferentially located near the base of the OHC (the locations would result in a change of the protocol discussed here). Further, it is desirable if the base is mechanically constrained in some way (e.g., perhaps held in Deiter’s cell), as then the displacement at the nearly collocated probe is small. The dynamic loading stiffness of the two experiments is denoted by 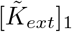 and 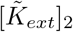. The measured NLC under these two boundary conditions is denoted as 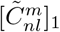 and 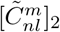. We have assumed that the nonlinear charge can be separated from the linear contribution during a whole-cell voltage controlled experiment [47]. The measured displacement-to-voltage transfer functions (or eM) are denoted by 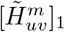 and 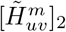. Using model Eqs. 3, 4, 16, and 17, an estimate of the electromechanical force coefficient, 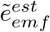 over the frequency range where measurements are made can be written as

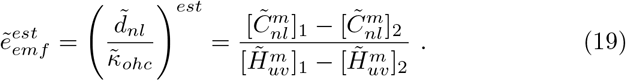

**Fig. 5.**
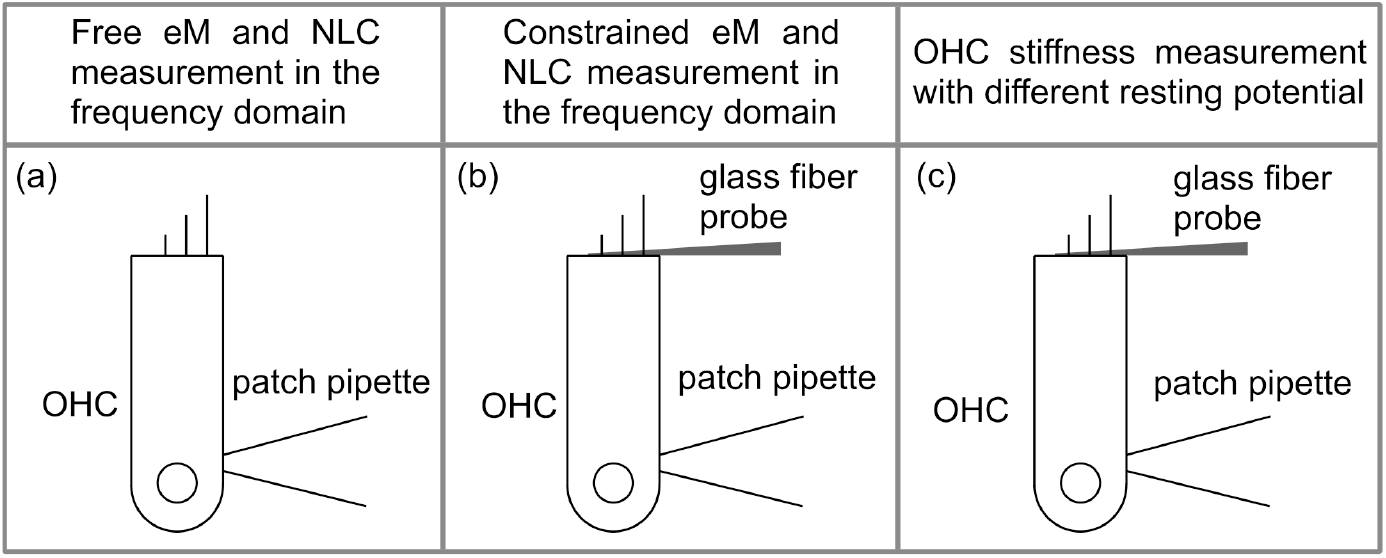
Schematics of the proposed experimental configurations using a patch pipette. eM and NLC measured (a) without and (b) with a glass fiber or atomic force microscope (AFM) probe as in 31, 52. (c) OHC stiffness measured under different resting potentials using the approach of 31.

The advantage of this approach is that 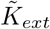does not enter into these relations. With the estimate of 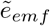 in hand, we can manipulate Eq. 4 to yield an estimate of the isometric NLC, 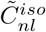, as

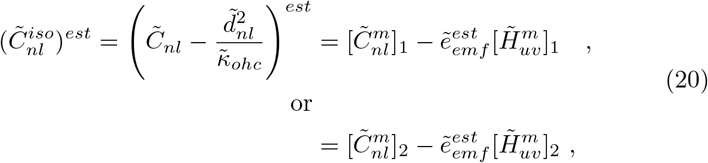

where two estimates of 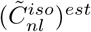are obtained.

With 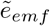and 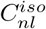 estimated, a procedure to estimate 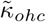 is needed. Previous OHC experiments have focused on obtaining the low frequency, quasistatic compliance of the cell or finding its stiffness at a few frequencies (e.g., 31, 53–55). But, to our knowledge, the full frequency dependence of the OHC dynamic stiffness, including rate dependent effects, has not been obtained. One approach is to first estimate the external load’s dynamic stiffness 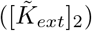 either through modeling or experiment [56], and extend the quasistatic approach put forth in 31 to the frequency domain. Identifying 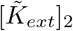 would be useful not only for estimating 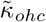 but also for providing a consistency test for our estimates of 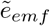 and 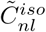. However, we propose an alternative approach by estimating the constituent model parameters of 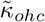: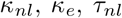, and *τ*_*e*_ (Eqs. 5 and 6). We describe one approach to achieving this next.

An experimental configuration described in 31 (schematically shown Fig. 5 (c)) could be used to estimate the OHC compliance at different resting potentials (the static stiffness of the probe is needed, as described in 31). Fig. 6(a) schematically shows a potential dependence of the OHC compliance on the resting potential. The compliance measured at very hyperpolarized or depolarized potentials provides an estimate of the elastic compliance *κ*_*e*_, as the gating compliance (*κ*_*nl*_) in the long or short state is taken as zero. The peak compliance in Fig. 6(a) provides an estimate of *κ*_*ohc*_ from which *κ*_*nl*_ is found from the difference between *κ*_*ohc*_ and *κ*_*e*_. An independent approach to estimate *κ*_*ohc*_ can be obtained by noticing that *κ*_*ohc*_ = *d*_*nl*_*/e*_*emf*_ (Eq. 8). We can use the low frequency estimates of *e*_*emf*_ (using Eq. 19) and *d*_*nl*_ (experimentally determined as the low frequency free eM of the cell, as shown in Fig. 5(a)). This technique serves as a consistency test for the OHC compliance near a resting potential.

**Fig. 6.**
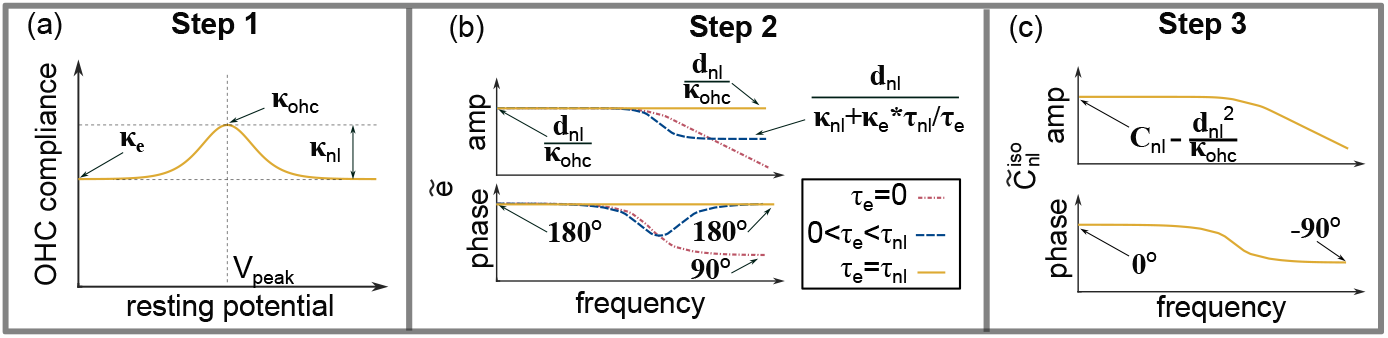
Proposed steps for parameter estimation of the 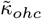. (a) Step 1: Estimate the dependence of the quasi-static compliance on the resting potential (following, e.g., 31, 53, 54). Values near *V*_*peak*_ and at highly depolarized or hyperpolarized values can be used to determine the value of *κ*_*e*_, *κ*_*nl*_, and *κ*_*ohc*_ (the full range of values are not needed). The cartoon of OHC compliance is in accordance with the two-state Boltzmann theory [31]. However, other nonlinear quasi-static constitutive theories are possible, and can be substituted for Boltzmann model [19, 36, 53]; (b) Step 2: Using the frequency dependence of the magnitude and phase of 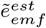 as estimated via Eq. 19 enables the ratio between *τ*_*nl*_ and *τ*_*e*_ to be determined by analysis of the high frequency asymptotes as exemplified for 0 *< τ*_*e*_ ≤ *τ*_*nl*_; (c) Step 3: undetermined parameter *τ*_*nl*_ can be estimated by curve fitting with the isometric NLC (amplitude and phase using Eq. 12).

Next, we discuss how to estimate the rate constants. A sketch of a prototypical frequency dependence of the magnitude and phase of 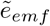 as shown in Fig 6(b) is used as the basis to estimate the ratio of two rates, *τ*_*nl*_*/τ*_*e*_. For instance, if the force factor exhibits a low-pass characteristic with the phase changing from 180 to 90 degrees (red dashed line), the value of *τ*_*e*_ is likely much smaller than *τ*_*nl*_. If the amplitude and phase hold constant through the entire frequency range, this implies that rates of the two contributions are similar in value (e.g., *τ*_*nl*_ = *τ*_*e*_, the gold line). In general, the ratio can be determined by fitting the Eq. 8 to the measurement (Fig. 6(b) blue curve). With the ratio, *τ*_*nl*_*/τ*_*e*_ now estimated, their individual values can then be determined by fitting the amplitude and phase of 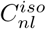 response simultaneously, as shown in Fig. 6(c).

High frequency whole-cell patch-clamp measurements would provide an incisive tool for resolving the debate about the effectiveness of the OHC at high frequencies by identifying rate dependent limits of OHC electromechanical behavior. The experiments proposed in Figs. 5-6 are similar to those presented in 22, 31, 54. However, we acknowledge there are a number of practical challenges facing these measurements. For instance, as documented in 31, the OHC may slip at the patch location while under mechanical load. This is problematic because the resting (static) mechanical load, voltage, and turgor pressure must be controlled in experiments in order to provide reliable estimates of the model parameters. This slipping is the reason for our recommendation that the cell be patched at the base which is held stationary. Also, technical challenges have limited the high-frequency measurement of OHC behavior due to the patch-clamp bandwidth [3]. Improved amplifier technology, such as that recently used for macropatch experiments [48]), holds the potential to increase the bandwidth for whole-cell measurements. The macropatch technique itself can access higher frequencies [11]). The NLC and eM results measured with this technique have shown a much lower cut-off frequency behavior [11, 21], although more recent results place the cut-off frequency at a higher value [48]. However, these experiments also have limitations due to the mechanical conditions. The patched portion of the membrane is loaded with a roughly 2-micron diameter fluid-filled pipet, which presents a viscous-(not mass-) dominated fluid impedance to the membrane, a feature that would itself introduce a rate dependence that would saturate with increasing fluid column height. Moreover, the mechanical boundary conditions of the patched membrane are not completely quantified as there is uncertainty with regard to both the membrane attachment to the cytoskeleton as well as the mechanical boundary conditions at the interface between the membrane and the suctioning pipet. For these reasons, we believe the whole-cell measurement may provide a more direct route to parameter estimation.

## 3 Discussion

In theories of cochlear amplification, based on OHC-somatic force generation, OHCs contribute to the sensitivity of the mammalian cochlea by amplifying traveling waves through electrical-to-mechanical energy conversion. Since the work of 22, OHC electromechanical processes have been considered sufficiently fast to contribute over the entire mammalian frequency range. However, other measurements, including some more recent data, indicate the cut-off frequency of the free eM and NLC of the OHC is much lower [11, 21, 47, 49] which motivates a re-examination of rate dependence in OHC mechanics. The quantities *κ*_*nl*_, *κ*_*e*_, *τ*_*nl*_, and *τ*_*e*_ are central to mechanistically predicting OHC behavior, but the experimental identification of these four parameters, except for *τ*_*nl*_, has not been extensively explored experimentally, and has been theoretically investigated only recently [19, 21]. We have highlighted that the frequency dependence of eM and NLC are not the same as that of electrical to mechanical force generation, as exemplified in Fig. 4. This means that models that fit *in-vitro* eM and NLC experimental results, can evince very different frequency dependencies of the electromechanical coupling factor depending on the ratios *κ*_*nl*_*/κ*_*e*_ and *τ*_*nl*_*/τ*_*e*_. Hence, solely focusing on the NLC and eM does not close the discussion of prestin-based amplification, because the frequency dependence of the electromechanical force coefficient, 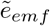, remains undetermined. This factor is seen to regulate mechanical power injection to the surrounding structures (Eq. 18). Further, we showed, perhaps for the first time, that the isometric NLC, 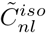 and not the free NLC, 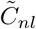, is most important in characterizing the transmembrane impedance (see Eqs. 4 and 12); the difference between these two quantities depends on the strength and frequency dependence of electromechanical coupling, which has yet to be completely described. Finally, there is the intriguing finding that if the NLC decreases at higher frequencies, then the lateral membrane presents an increasing impedance to MET currents (see Eq. 11), a feature that could impact electrical-to-mechanical force production. Overall, even if only a subset of our proposed measurements are possible (like those to determine the frequency dependence of 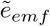 and 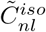, as presented in Eqs. 19-20), it would represent a crucial step forward in determining the function of the OHC over the audio frequency range, hopefully moving us closer to settling this important debate over the effectiveness of OHC force generation at high frequencies.

## 4 Methods

This section consists of two parts. In the first, we develop the nonlinear, rate-dependent constitutive equations governing the OHC response. We next perform a consistent linearizataion of these equations as used in the main text (Eqs.1 and 2). In the second part, we show how we estimate the model parameters to fit the experimental data.

### Linearized governing equations for OHCs

The OHC is a fluid-filled cell of nearly cylindrical shape *in situ* when carefully extracted from the organ of Corti [3, 31]. Only the axial deformation of the OHC is considered here. The OHC is modeled as a prestin-based motile element in series with a viscoelastic element. In Fig. 7(a), the nonlinear mechanics of the prestin related conformal change is represented. The viscoelastic element is shown in Fig. 7(b)) represents the stiffness and damping effect of the lipid bilayer which is connected in series with the motor element.

**Fig. 7.**
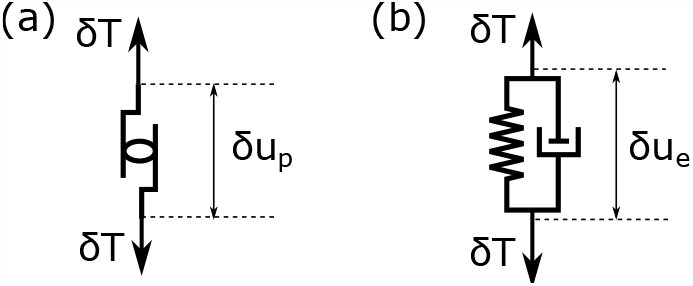
Free body diagram of (a) the electromotile element (which depends nonlinearly on *δT* and *δV*) and (b) viscoelastic element undergoing small perturbations from the resting state.

### Rate-dependent model for the electromotile element using two-state Boltzmann distribution

First, we assume that the electromotile element (or motor) in Fig. 7(a) possesses two conformal states, a long and a short state [31, 57], which can be characterized by conformal statistics defined by, 𝒫 (*T, V*) according to a two-state Boltzmann distribution. We will linearize this model about an equilibrium point to characterize the small signal response.

At equilibrium, the OHC is loaded by the force, *T*_0_, and voltage, *V*_0_, with a probability 𝒫 _0_. After the small perturbation *δT* and *δV* are introduced to the system, the linearized conformal probability becomes the summation of 𝒫 _0_ and the deviation from equilibrium, *δ*𝒫 as

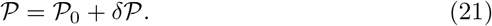

The perturbations of the displacement, *δu*_*p*_, and the nonlinear charge, *δQ*_*nl*_, from equilibrium caused by the disturbance are then described as

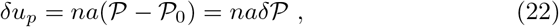

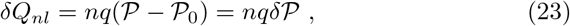

where *n* is the total number of electromotile proteins in the lateral wall of the OHC, *a* and *q* are the length change and charge transfer [58]. Here we assume that the nonlinear charge and nonlinear displacements are proportional to one another.

To consider the conformal state transition rate influences, the probability function 𝒫 is governed by Eq. 24 based on the Eyring equation [21, 59, 60] as

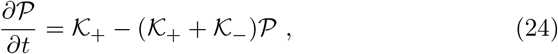

where the rate coefficients are defined by

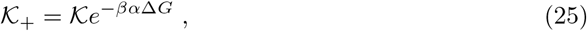

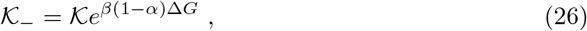

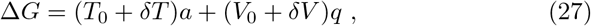

In Eqs. 25 and 26, 𝒦 relates to the energy barrier between the two states at the equilibrium, which is assumed as a constant [60]; 𝒦_+_ and 𝒦_−_ are the transition rate coefficients between the long and short state; the deviation of the constant *α* (0 ≤ *α* ≤ 1) from 1*/*2 indicates the asymmetry of the free energy at the two states; *β* = 1*/*(*k*_*B*_Θ), where *k*_*B*_ is Boltzmann’s constant and Θ is the temperature in Kelvin; ∆*G* represents Gibb’s free energy difference between the short and long states.

According to Eq. 24, the equilibrium probability at the condition of *T*_0_ and *V*_0_ is determined by

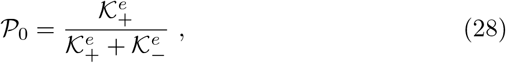

where 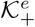and 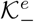 are the rate coefficients at *T*_0_ and *V*_0_ with energy difference ∆*G*_*e*_ = *T*_0_*a* + *V*_0_*q*.

The deviation of the probability 𝒫 from the equilibrium point 𝒫 _0_ caused by the small perturbations *δT* and *δV* can be described using the linear perturbation as:

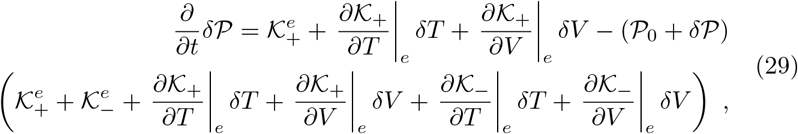

where | _*e*_ indicates the partial derivative is evaluated at the equilibrium condition.

Therefore, based on Eqs. 25-28, the linearized governing equation for *δ*𝒫 in Eq. 29 is further reduced to

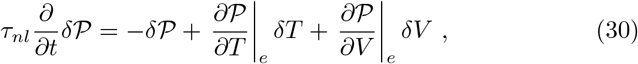

where

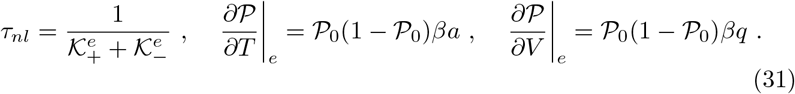

In the absolute-rate Eyring equation model, *τ*_*nl*_ attains its maximum value when the motor is at the half-probability state and decreases about that point. According to the relations between the displacement, nonlinear charge, and probability in Eqs. 22 and 23, the governing equations for the electromotile displacement *δu*_*p*_ and nonlinear charge *δQ*_*nl*_ are

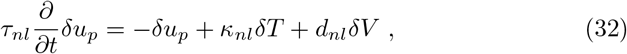

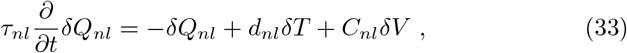

where

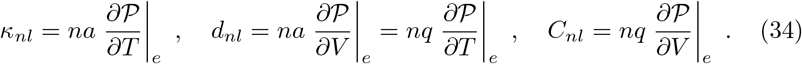

Finally, we note here that 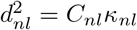 and 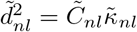.

### Rate-dependent model for the electromotile element using classical continuum mechanics constitutive theory

The governing equations shown in Eqs. 32 and 33 are derived from the 2-state Boltzmann model. On the other hand, we can employ the methods from continuum mechanics [61] to arrive at the coupled, linearized equations for quasi-static perturbations about equilibrium as:

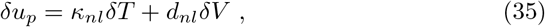

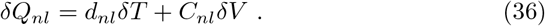

The parameters in Eqs. 35 and 36 can be derived from the free energy *W* (*T, V*) of the OHC [61], which are

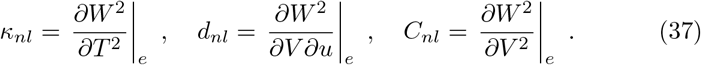

Non-equilibrium transitions are modeled by introducing rate-dependent terms (proportional to *τ*_1_ and *τ*_2_) in the following manner [19, 61]:

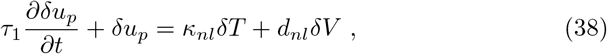

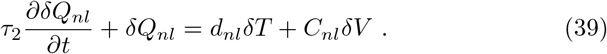

In Eqs. 32 and 33, the rate dependence *τ*_*nl*_ is derived for both the displacement and charge. However, the losses in the mechanical and electrical domains may be different, resulting in distinct numerical values for the rate *τ*_1_ and *τ*_2_. When *τ*_1_ = *τ*_2_ = *τ*_*nl*_, the governing equations derived from the classic mechanical relations are the same as Eqs. 32 and 33 from the Eyring rate equation, which demonstrates the consistency in modeling the OHC from both approaches. In the study, the same rate *τ*_1_ = *τ*_2_ assumption is adopted throughout.

In the Boltzmann theory presented above, we have assumed that the non-linear electromotile strain *u*_*p*_ is linearly proportion to the nonlinear charge transfer *Q*_*p*_ in Eqs. 22-23. If we make the same assumption in the thermodynamic model, this requires that the fraction 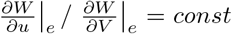 and that ^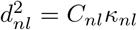^, as in the Boltzmann-based theory.

### Mechanical model for the viscoelastic model

Because prestin is intercalated in the lipid bilayer of the plasma membrane [32–34], active and passive prestin forces are transmitted through the lipid portion of the membrane bilayer. Therefore, we include a viscoelastic contribution in series with the motor element as shown in Fig. 7(b). According to Fig. 7(b), the rate-dependent equation of this contribution is given by

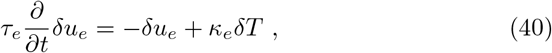

where *κ*_*e*_ is the compliance of the lipid bilayer, and *τ*_*e*_ is the rate dependence of the viscous element. We assume a simple integer-order damping for this viscous force, rather than the fractional calculus that is sometimes used for losses in biological tissue [19]. This viscous element represents a rate dependence associated with the deformation of the lipid bilayer in the plasma membrane potentially different from the rate dependence of any conformal changes in prestin.

### Linearized OHC dynamic model

As demonstrated in the Appendix of 19, the homogenized contributions of all the individual motor protein-lipid bilayer components can additively be decomposed so that the total formation associated with the perturbation is given by *δu* = *δu*_*p*_ + *δu*_*e*_. Finally, the total perturbed charge across the cell membrane is given by *δQ* = *δQ*_*nl*_ +*δQ*_*lin*_ where the linear contribution to the total current is often carefully identified and subtracted from experimentally presented results (e.g., 39), and we will follow that convention.

Eqs. 32-40 govern the time domain response of the OHC to mechanical and electrical excitation. In order to simplify the analysis, we will transform from the time domain to the frequency domain to obtain the frequency response of the system by substituting a time-harmonic dependence of the form *e*^*jωt*^ for each variable, e.g., setting *δu*_*p*_ to *δũ*_*p*_*e*^*jωt*^ and eliminate *δũ*_*e*_ and *δũ*_*p*_ in favor of directly measurable total cellular deformation using *δũ* = *δũ*_*p*_ + *δũ*_*e*_. This transformation is applied to Eqs. 32-40 to obtain

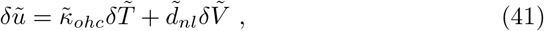

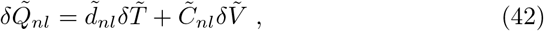

where,

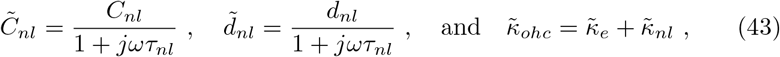

with

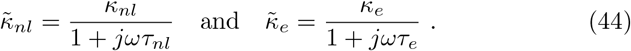

The relations may be converted to a force-charge form as

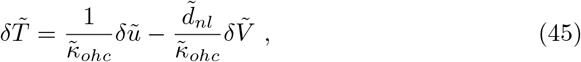

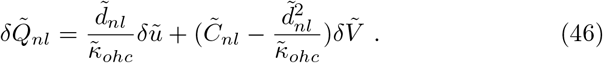

With the relations presented in Eqs. 45 and 46, the model of the OHC connecting to an external load can be further established.

### Parameter Estimation for OHCs

We next describe how the six intrinsic model parameters (*κ*_*nl*_, *κ*_*e*_, *τ*_*nl*_, *τ*_*e*_, *C*_*nl*_, and *d*_*nl*_) and the three external loading quantities (*K*_*ext*_, *η*_*ext*_, and *M*_*ext*_) as recorded in Table 1 were determined.

For OHC 65 and OHC 84, the intrinsic parameters *κ*_*ohc*_ and *d*_*nl*_ can be directly determined by the experimental data in 22. According to Frank’s measurement, the OHC compliance *κ*_*ohc*_ is linearly dependent on the OHC length, *L*_*ohc*_, as *κ*_*ohc*_ = *C* · *L*_*ohc*_. To better fit the experimental data in Fig. 3, the coefficient *C* is chosen to be 2.28 *μN* ^−1^ for OHC 65 and 3.10 *μN* ^−1^ for OHC 84. The OHC length is 55.6 *μm* for OHC 65 and 24 *μm* for OHC 84. Based on Eq. 16, the value of *d*_*nl*_ can be estimated by the load-free eM at static or quasi-static conditions. The value of *d*_*nl*_ in Table 1 is approximated by the eM amplitude at 100 Hz in Fig. 3(a). Since there is no electrical data documented for OHC 65 and OHC 84, the parameter *C*_*nl*_ is left undetermined. To estimate *κ*_*nl*_, Eqs. 31 and 34 are utilized. The values of the length change *a* and charge transfer *q* are taken from the literature to be 64 *fm* [41] and 0.72*e* [62]. The number of proteins *n* is calculated by the motile density 9 *×* 10^3^ *μm*^−2^ multiplying the cell area (= 2*πr* · *L*_*ohc*_, where *r* = 5*μm*) [62]. With the parameters of *a, q, n*, and *d*_*nl*_ determined above, the equilibrium probability is estimated to be 0.125 for OHC 65 and 0.05 for OHC 84, supporting that the holding potential in Frank’s measurement is far away from *V*_*peak*_ [21]. The compliance *κ*_*nl*_ in the model is then calculated to be 1.7 *m/N* for OHC 65 and 0.32 *m/N* for OHC 84. The elastic compliance *κ*_*e*_ is estimated by *κ*_*ohc*_ − *κ*_*nl*_.

For the external loading quantities used to fit the measurements of OHC 65 and OHC 84, 22 estimated the external mass load *m*_*ext*_ as around 6% of the cell mass (cell mass is calculated by *πr*^2^*L*_*ohc*_*ρ*, where *ρ* is the density of water) in order to fit their data. In Table 1, for OHC 65 and OHC 84, we found a value of 5% of the cell mass provided a slightly better to fit the experiments, and used that value.The external damping *η*_*ext*_ is determined by the quality factor demonstrated in 22. For the eM responses in Figs. 3(a) and (b), the external stiffness *K*_*ext*_ is zero. Whereas the electromechanical coupling responses in Figs. 3(c) and (d) simulate the loading case as *K*_*ext*_ approaches infinity. With the estimation of other parameters, the rate *τ*_*nl*_ is determined by fitting the experimental curves.

For OHC 1, based on the relation between the linear capacitance and the OHC length, the length of OHC 1 is estimated to be 78 *μm* for a linear capacitance of 22.5 *pF* [3]. As the data in Fig. 4 is normalized, Eqs. 31 and 34 are employed to estimate the value of *d*_*nl*_, *C*_*nl*_, and *κ*_*nl*_. Since the holding potential of the OHC 1 measurement is reported around *V*_*peak*_, the equilibrium probability is set to be 0.5. The charge transfer *q* is varied to be 0.9*e* to better match the NLC value of 28 *pF* found in 47. The length change *a* keeps at 64 *fm*. The number of proteins *n* is calculated with the motile density 9 *×* 10^3^ *μm*^−2^ as described for OHC 65 and OHC 84. After the estimation of *d*_*nl*_, *C*_*nl*_, and *κ*_*nl*_, the OHC compliance *κ*_*ohc*_ is then determined to be 123 *m/N* according to Eq. 8 based on the experimental evidence that the static electromechanical force coefficient *e*_*emf*_ holds constant around -0.1 *nN/mV* along the cochlea [31]. The elastic compliance *κ*_*e*_ can then be likewise determined by *κ*_*ohc*_ − *κ*_*nl*_. Since the external added mass *m*_*ext*_ does not show great influence on responses below 6 *kHz, m*_*ext*_ for OHC 1 is estimated as 8*ρr*^3^*/*3 using a simple entrained mass approximation, where *ρ* is the water density, *r* is the cell radius [63]. Based on the current experimental and theoretical evidence, the value of *τ*_*nl*_ shows a clear relation with the cut-off frequency of the measured NLC. Therefore, for OHC 1, *τ*_*nl*_ is estimated by the cut-off frequency of the measured NLC, and *η*_*ext*_ is then determined by fitting the responses.

Regarding the parameter estimation described above, there is insufficient data to estimate the value of rate *τ*_*e*_. In the paper, *τ*_*e*_ is varied in the range of 0 to *τ*_*nl*_ to investigate the influence on the responses.

## Acknowledgements

This work was supported by NIH-NIDCD R01DC04184. The content is solely the responsibility of the authors and does not necessarily represent the official views of the National Institutes of Health.

## Author contributions statement

W.C. and K.G. developed the theory and analyzed the results. W.C. and K.G. wrote and reviewed the manuscript. K.G. initialized the research.

## References

[1] Fettiplace, R.: Hair cell transduction, tuning, and synaptic transmission in the mammalian cochlea. Compr. Physiol. 7(4), 1197–1227 (2017)

[2] Johnson, S.L., Beurg, M., Marcotti, W., Fettiplace, R.: Prestin-driven cochlear amplification is not limited by the outer hair cell membrane time constant. Neuron. 70(6), 1143–1154 (2011)

[3] Ashmore, J.F.: Cochlear outer hair cell motility. Physiol. Rev.l. 88, 173–210 (2008)

[4] Ashmore, J., Avan, P., Brownell, W.E., Dallos, P., Dierkes, K., Fettiplace, R., Grosh, K., Hackney, C.M., Hudspeth, A.J., Jülicher, F., Lindner, B., Martin, P., Meaud, J., Petit, C., Santos Sacchi, J.R., Canlon, B.: The remarkable cochlear amplifier. Hearing Research 266(1), 1–17 (2010). 10.1016/j.heares.2010.05.001. Special Issue: Annual Reviews 2010

[5] Fisher, J.A., Nin, F., Reichenbach, T., Uthaiah, R.C., Hudspeth, A.J.: The spatial pattern of cochlear amplification. Neuron 76(5), 989–997 (2012)

[6] Corey, D.P., Akyuz, N., Holt, J.P.: Function and dysfunction of tmc channels in inner ear hair cells. Cold Spring Harb. Perspect. Med. 9(10) (2019)

[7] Dallos, P., Wu, X., Cheatham, M.A., Gao, J., Zheng, J., Anderson, C.T., Jia, S., Wang, X., Cheng, W.H., Sengupta, S., He, D.Z.: Prestin-based outer hair cell motility is necessary for mammalian cochlear amplification. Neuron 3(58), 333–339 (2008)

[8] Brownell, W.E., Bader, C.R., Bertrand, D., Ribaupierre, Y.D.: Evoked mechanical responses of isolated cochlear hair cells. Science. 227, 194–196 (1985)

[9] Dewey, J.B., Altoe, A., Shera, C.A., Applegate, B.E., Oghalai, J.S.: Cochlear outer hair cell electromotility enhances organ of corti motion on a cycle-by-cycle basis at high frequencies in vivo. Proc. Natl. Acad. Sci. USA. 118(43) (2021)

[10] Sasmal, A., Grosh, K.: Unified cochlear model for low-and highfrequency mammalian hearing. Proc. Natl. Acad. Sci. U.S.A. 116(28), 13983–13988 (2019)

[11] Gale, J.E., Ashmore, J.F.: The outer hair cell motor in membrane patches. Pflügers. Arch. Eur. J. Physiol. 434, 267–271 (1997)

[12] Santos-Sacchi, J.: On the frequency limit and phase of outer hair cell motility: effects of the membrane filter. J. Neurosci. 12(5), 1906–1916 (1992)

[13] Altoè, A., Shera, C.A.: The long outer-hair-cell rc time constant: A feature, not a bug, of the mammalian cochlea. J. Assoc. Res. Otolaryngol. 24(2), 129–145 (2023)

[14] Ramamoorthy, S., Deo, N.V., Grosh, K.: A mechano-electro-acoustical model for the cochlea: response to acoustic stimuli. J. Acoust. Soc. Am. 121(5), 2758–2773 (2007)

[15] Housley, G.D., Ashmore, J.F.: Ionic currents of outer hair cells isolated from the guinea-pig cochlea. J. Physiol. 448(1), 73–98 (1992)

[16] van der Heijden, M., Vavakou, A.: Rectifying and sluggish: outer hair cells as regulators rather than amplifiers. Hear. Res. 423, 108367 (2022)

[17] Vavakou, A., Cooper, N.P., van der Heijden, M.: The frequency limit of outer hair cell motility measured in vivo. Elife. 8, 47667 (2019)

[18] Altoè, A., Charaziak, K.K., Shera, C.A.: Dynamics of cochlear nonlinearity: Automatic gain control or instantaneous damping? J. Acoust. Soc. Am. 142(6), 3510–3519 (2017)

[19] Rabbitt, R.D.: The cochlear outer hair cell speed paradox. Proc. Natl. Acad. Sci. USA. 117(36), 21880–21888 (2020)

[20] Gale, J.E., Ashmore, J.F.: Charge displacement induced by rapid stretch in the basolateral membrane of the guinea-pig outer hair cell. Proc. Royal Soc. (London) B, Biol. Sci. 255, 233–249 (1994)

[21] Santos-Sacchi, J., Iwasa, K.H., Tan, W.: Outer hair cell electromotility is low-pass filtered relative to the molecular conformational changes that produce nonlinear capacitance. J. Gen. Physiol. 151(12), 1369–1385 (2019)

[22] Frank, G., Hemmert, W., Gummer, A.: Limiting dynamics of highfrequency electromechanical transduction of outer hair cells. Proc. Natl. Acad. Sci. USA. 96, 4420–4425 (1999)

[23] Scherer, M.P., Gummer, A.W.: Vibration pattern of the organ of corti up to 50 khz: evidence for resonant electromechanical force. Proc. Natl. Acad. Sci. U.S.A. 101(51), 17652–17657 (2004)

[24] Grosh, K., Zheng, J.F., Zou, Y., de Boer, E., Nuttall, A.L.: Highfrequency electromotile responses in the cochlea. J. Acoust. Soc. Am. 115(5), 2178–2184 (2004)

[25] Ren, T., He, W., Kemp, D.: Reticular lamina and basilar membrane vibrations in living mouse cochleae. Proc. Natl. Acad. Sci. U.S.A. 113(35), 9910–9915 (2016)

[26] Nankali, A., Shera, C.A., Applegate, B.E., Oghalai, J.S.: Interplay between traveling wave propagation and amplification at the apex of the mouse cochlea. Biophys. J. 121(15), 2940–2951 (2022)

[27] Frost, B., Olson, E.S.: Model of cochlear microphonic explores the tuning and magnitude of hair cell transduction current. Biophys. J. 120(17), 3550–3565 (2021)

[28] Nam, J.H., Fettiplace, R.: Force transmission in the organ of corti micromachine. Biophys. J. 98(12), 2813–2821 (2010)

[29] Dewey, J.B., Applegate, B.E., Oghalai, J.S.: Amplification and suppression of traveling waves along the mouse organ of corti: evidence for spatial variation in the longitudinal coupling of outer hair cell-generated forces. J. Neurosci. 39(10), 1805–1816 (2019)

[30] Zhou, W., Nam, J.H.: Probing hair cell’s mechano-transduction using two-tone suppression measurements. Sci. Rep. 9(1), 4626 (2019)

[31] Iwasa, K.H., Adachi, M.: Force generation in the outer hair cell of the cochlea. Biophys. J. 73, 546–555 (1997)

[32] Dehghani-Ghahnaviyeh, S., Zhao, Z., Tajkhorshid, E.: Lipid-mediated prestin organization in outer hair cell membranes and its implications in sound amplification. Nat. Commun. 13(1), 6877 (2022)

[33] Bavi, N., Clark, M.D., Contreras, G.F., Shen, R., Reddy, B., Milewski, W., Perozo, E.: Cryo-em structures of prestin and the molecular basis of outer hair cell electromotility. bioRxiv., 2021–08 (2021)

[34] Holley, M.C., Ashmore, J.F.: A cytoskeletal spring in cochlear outer hair cells. Nature. 335(6191), 635–637 (1988)

[35] Tolomeo, J.A., Steele, C.R., Holley, M.C.: Mechanical properties of the lateral cortex of mammalian auditory outer hair cells. Biophys. J. 71(1), 421–429 (1996)

[36] Deo, N., Grosh, K.: Two-state model for outer hair cell stiffness and motility. Biophys. J. 86(6), 3519–3528 (2004)

[37] Adachi, M., Iwasa, K.H.: Electrically driven motor in the outer hair cell: Effect of a mechanical constraint. Proc. Natl. Acad. Sci. USA. 96, 7244–7249 (1999)

[38] Deo, N., Grosh, K.: Simplified nonlinear outer hair cell models. J. Acoust. Soc. Am 117(4), 2141–2146 (2005)

[39] Santos-Sacchi, J., Tan, W.: The frequency response of outer hair cell voltage-dependent motility is limited by kinetics of prestin. J. Neurosci. 38(24), 5495–5506 (2018)

[40] Levic, S., Lukashkina, V.A., Simoes, P., Lukashkin, A.N., Russell, I.J.: A gap-junction mutation reveals that outer hair cell extracellular receptor potentials drive high-frequency cochlear amplification. J. Neurosci 42(42), 7875–7884 (2022)

[41] Iwasa, K.H.: Energy output from a single outer hair cell. Biophys. J. 111(11), 2500–2511 (2016)

[42] Dimitriadis, E.K., Chadwick, R.S.: Solution of the inverse problem for a linear cochlear model: A tonotopic cochlear amplifier. J. Acoust. Soc. Am 106(4), 1880–1892 (1999)

[43] Corbitt, C., Farinelli, F., Brownell, W.E., Farrell, B.: Tonotopic relationships reveal the charge density varies along the lateral wall of outer hair cells. Biophys. J. 102(12), 2715–2724 (2012)

[44] Ashmore, J.F.: A fast motile response in guinea-pig outer hair cells: the cellular basis of the cochlear amplifier. J. Physiol. (Lond.). 388, 323–347 (1987)

[45] Iwasa, K.H.: Effect of membrane motor on the axial stiffness of the cochlear outer hair cell. J. Acoust. Soc. Am. 107(5), 2764–2766 (2000)

[46] Tolomeo, J.A., Steele, C.R.: Orthotropic piezolelectric properties of the cochlear outer hair cell wall. J. Acoust. Soc. Am. 97(5.1), 3006–3011 (1995)

[47] Santos-Sacchi, J., Tan, W.: Coupling between outer hair cell electromotility and prestin sensor charge depends on voltage operating point. Hear. Res. (2022)

[48] Santos-Sacchi, J., Bai, J.P., Navaratnam, D.: Megahertz sampling of prestin (slc26a5) voltage-sensor charge movements in outer hair cell membranes reveals ultrasonic activity that may support electromotility and cochlear amplification. J. Neurosci. (2023)

[49] Santos-Sacchi, J.: The speed limit of outer hair cell electromechanical activity. Hno (3), 159–164 (2019)

[50] Meaud, J., Grosh, K.: Coupling active hair bundle mechanics, fast adaptation, and somatic motility in a cochlear model. Biophys. J. 100(11), 2576–2585 (2011)

[51] Wang, Y., Steele, C.R., Puria, S.: Cochlear outer-hair-cell power generation and viscous fluid loss. Sci. Rep. 6(1), 19475 (2016)

[52] Hallworth, R.: Passive compliance and active force generation in the guinea pig outer hair cell. J. Neurophysiol. 74(6), 2319–2328 (1995)

[53] He, D.Z.Z., Dallos, P.: Somatic stiffness of cochlear outer hair cells is voltage-dependent. Proc. Natl. Acad. Sci. USA. 96(14), 8223–8228 (1999)

[54] Hallworth, R.: Absence of voltage-dependent compliance in highfrequency cochlear outer hair cells. J. Assoc. Res. Oto. 8(4), 464–473 (2007)

[55] He, D.Z.Z., Dallos, P.: Properties of voltage-dependant somatic stiffness of cochlear outer hair cells. J. Assoc. Res. Otolaryngol. 01, 64–81 (2000)

[56] Howard, J., Clark, R.L.: Mechanics of Motor Proteins and the Cytoskeleton. Sinauer Associates, Sunderland, Massachusetts (2002)

[57] Iwasa, K.H.: A membrane motor model for the fast motility of the outer hair cell. J. Acoust. Soc. Am. 96(4), 2216–2224 (1994)

[58] Iwasa, K.H.: Effect of stress on the membrane capacitance of the auditory outer hair cell. Biophys. J. 65, 492–498 (1993)

[59] Choe, Y., Magnasco, M.O., Hudspeth, A.J.: A model for amplification of hair-bundle motion by cyclical binding of ca2+ to mechanoelectricaltransduction channels. Proc. Natl. Acad. Sci. USA. 95(26), 15321–15326 (1998)

[60] Keener, J., Sneyd, J.: Mathematical Physiology: II: Systems Physiology. NY: Springer New York, New York (2009)

[61] Anand, L., Govindjee, S.: Continuum Mechanics of Solids. Oxford University, Oxford (2020)

[62] Iwasa, K.H.: A two-state piezoelectric model for outer hair cell motility. Biophys. J. 81, 2495–2506 (2001)

[63] Krishnaswami, G.S., Phatak, S.: The added mass effect and the higgs mechanism. Reson. 25(2), 191–213 (2020)

